# Chitosan nanoparticle vaccine loaded with crude extracellular proteins of *C. perfringens* and *Salmonella* flagella is partially protective against necrotic enteritis in broiler chickens

**DOI:** 10.1101/2020.10.15.340661

**Authors:** Gabriel Akerele, Nour Ramadan, Muhammed Mortada, Revathi Shanmugasundaram, Sankar Renu, Gourapura J. Renukaradhya, Ramesh K Selvaraj

## Abstract

Necrotic enteritis (NE) causes significant economic losses and food shortages world-wide. There are currently no licensed commercial vaccines against NE in broilers. Chitosan nanoparticles were formulated with extracellular proteins of *C. perfringens* surface-tagged with *Salmonella* flagellar proteins. One-day-old male broiler chicks were completely randomized to 3 treatments: Non-vaccinated non-challenged as negative control, Vaccinated-challenged, and non-vaccinated challenge as positive control. On day of hatch, d7, and d14 post-hatch, vaccinated-challenged birds were orally gavage with 50μg vaccine in 0.5ml PBS while positive control birds were gavage with 0.5ml PBS only. Birds in the vaccinated-challenged and positive control groups were orally infected on d14 post-hatch, with 5,000 oocysts/bird of *E. maxima*, followed by log 8 CFU of a virulent strain of C. perfringens on d19, d20, and d21 post-hatch. From d14 to 21 and d14 to 28 post-hatches, mortality in the vaccinated-challenged group was higher than that in the positive control group, approaching statistical significance (p=0.07). On d21 post-hatch, the mean lesion score of 3 birds/cage in the vaccinated-challenged group was higher than the positive control group, approaching statistical significance (p = 0.05). From d 14 to 28 post-hatch, the feed intake was higher and feed conversion ratio lower in the vaccinated-challenged group compared to the positive control group (p<0.05). On d21 post-hatch, antigen specific recall proliferation in the vaccinated-challenged group was higher than that in the negative and positive control groups (p<0.05). On d21 post-hatch, cecal tonsils CD8+ T lymphocytes expression in the vaccinated-challenged group was similar to the negative control group (p>0.05) but higher than that in the positive control group (p<0.05). Finally, vaccination resulted in an increase in ileal mRNA levels of zonula occluding on d21 post-hatch. In conclusion, there were numerical but not statistically significant decrease in NE lesions and mortality in vaccinated and challenged broilers. Further studies are needed to improve the efficacy of the vaccine and understand the mechanism underlying protection in vaccinated birds.

## Introduction

Necrotic enteritis (NE) is a disease of *Clostridium perfringens* that affects poultry gut health. Losses are estimated to be at least 2 billion USD annually from clinical and subclinical infections. The resurgence in infection is due to the voluntary or regulatory withdrawal in the use of antibiotics as growth promoters [1]. Poultry NE is a multifactorial disease with a complex etiology. However, the overgrowth of commensal or virulent *C. perfringens* strains is the primary cause of infection [2]. Current intervention strategies include the use of feed additives, improving biosecurity/management practices, anticoccidial drugs and vaccinations [3, 4]. There are currently no commercial vaccines against NE in broilers. Most commercial vaccine products are anti coccidia vaccines because coccidiosis is one of the most important predisposing conditions for NE [5].

Two important considerations for any vaccine design strategy are antigen selection and route of delivery. Subunit antigens are generally safer than live antigens for vaccines. Protein subunits in the extracellular secretion of *C. perfringens* (ECP) have been explored as antigens for candidate vaccines against NE with mixed results, with the most promising results involving parenteral routes of administration [6, 7, 8]. Parenteral antigen delivery is neither economical nor practical for large commercial operations. However, unencapsulated, subunit antigens such as proteins are denatured or broken down by the low pH conditions and gut enzymes when administered by oral gavage. Unencapsulated, subunit antigens are therefore inefficiently delivery to inductive sites, compromising their immunogenicity and protective efficacy [9]. Therefore, there is a need to develop immunogenic, safe and efficacious vaccines based on *C. perfringens* ECP that can be administered by orally to help mitigate production losses to NE in the poultry industry.

The safety and immunogenicity of nanoparticles as encapsulating and antigen delivery vehicles have been well documented [10, 11, 12]. Chitosan has emerged as a promising candidate for effective antigen delivery [13, 14]. A previous study conducted in our laboratory indicates that chitosan nanoparticles entrapped with *C. perfringens* ECP and surface-tagged with *Salmonella* flagella proteins is safe for oral vaccination and immunogenic in broiler chickens [15]. However, immunogenicity does not always correlate with protection. Therefore, further studies are required to characterize the protective efficacy of the previously synthesized chitosan nanoparticle vaccine against NE in broiler chickens.

The specific objectives of this study were to determine the immunogenicity and protective efficacy of an oral chitosan nanoparticle vaccine against experimentally induced NE. It is hypothesized that chitosan loaded with extracellular proteins of *C. perfringens* and delivered orally to broilers will stimulate antigen-specific immune responses and will reduce the severity of NE by improving boiler production performance, decreasing gut lesions.

## Materials and methods

### Experimental animals

All animal protocols were approved by the Institutional Animal Care and Use Committee of the Southern Poultry Research Group. Birds were monitored daily by trained veterinary care staff for diarrhea, bloody feces, lethargy, ruffled feathers, refusal to eat and dehydration in the course of the experiment. Birds that could not move or refused to eat were immediately humanely euthanized by carbon dioxide. A total of 2 animals reached the humane end point during the course of this study. All remaining birds were also humanely euthanized by carbon dioxide on the last day of sampling (day 28).

A total of 144 one-day-old male broiler chicks obtained from a commercial hatchery were raised in Petersime battery cages for 28 days at the Southern Poultry Research facility (Athens, GA). Chicks were weighed by pen and then randomly distributed to one of three treatments groups in a completely randomized design: Negative control group consisted of chicks that were not vaccinated and not challenged. The vaccinated-challenge group consisted of chicks that were both vaccinated and challenged. The positive control group consisted of chicks that were mock vaccinated and challenged. A cage was treated as a replicate. Each treatment was replicated in six cages (n = 6) of eight chicks per cage. Birds were raised under standard management practices and had *ad libitum* access to food and water. Feed intake, body weight and mortality were measured weekly from day of hatch. Mortality figures were compiled from birds that met the criteria for euthanasia as well as birds that died suddenly without meeting the criteria for euthanasia, after confirming NE by necropsy.

### Preparation and administration of vaccine, necrotic enteritis challenge and lesion scoring

A field strain of *C. perfringens* (a gift from Dr. C. Hofacre, Southern Poultry Research Group) was obtained. Chitosan nanoparticle vaccine was synthesized with native extracellular proteins of a virulent field strain of *C. perfringens* and surface tagged with flagella proteins according to methods detailed in a previous publication [15]. Synthesized nanoparticles were resuspended to mg/ml loaded protein in PBS at pH 7.2. On d 0 (day of hatch), d 4 and d 14 post-hatch, each bird in the vaccinated-challenged and positive control group was orally gavage with 50 μg loaded protein in nanoparticle suspension in 0.5 ml PBS and 0.5 ml PBS only, respectively.

Each bird in the vaccinated-challenged and positive control group was infected by oral gavage with 5,000 oocysts of *E. maxima* on d 14 post-hatch and then orally gavage with 1.0 × 10^8^ colony forming units of the same isolate of *C. perfringens* used to prepare the vaccine on d 19, 20 and 21 post-hatches.

Three randomly selected birds per pen were humanely euthanized by cervical dislocation and scored for NE lesions on d 21 post-hatch. The scoring system was based on a score of 0-3 as follows: 0 for a normal intestine, 1 for the presence of a slight mucus covering and loss of tone, 2 for severe NE and 3 for severe NE with the presence of blood in the lumen.

### Ex vivo recall response of splenic and cecal tonsil mononuclear cells of chickens

Cecal tonsil and spleen samples were collected from one bird per cage on d 18, and two birds per cage on d 21 and d 28 post-hatch after cervical dislocation. Single cell suspensions of cecal tonsils and spleen mononuclear cells were obtained as described earlier [16].

Approximately 5×10^4^ mononuclear cells/well were plated in duplicates per sample in 100μl of RPMI-1640 culture media (Sigma Aldrich, St. Louis, MO) supplemented with 10 % fetal bovine serum and 1 % Penicillin and Streptomycin. 100μl of RPMI-1640 culture media only (negative control) or 25μg, or 50 μg or 100μg of native ECP or 0.1 μg of Conclanavin A (Con A) in 100 μl of RPMI-1640 culture media was added to each well and incubated for 5 d at 37 °C in the presence of 5 % CO_2_. Lymphocyte proliferation was measured using [3-(4,5-dimethylthiazol-2-yl)-2,5-diphenyltetrazolium bromide] (MTT) assay as described earlier [13]. Values are reported as Optical density measured at 570 nm using a spectrophotometer.

### CD4^+^ and CD8^+^ cell percentages in cecal tonsils of vaccinated chickens

On d 18, 21 and 28 post-hatches, single cell suspensions of cecal tonsils and spleen mononuclear cells were obtained as described earlier [16]. The cell suspensions were concentrated for lymphocytes by density centrifugation over 1.077 g/mL histopaque (Sigma–Aldrich, St. Louis, MO). For CD4^+^ and CD8^+^ T cell analysis, single-cell suspensions of the cecal tonsils (1 × 10^6^ cells) were incubated with FITC-conjugated mouse anti-chicken CD4, PE-conjugated mouse anti-chicken CD8 (Southern Biotech, Birmingham, AL) at 1:200 dilution, and unlabeled mouse IgG at 1:200 dilution in a 96-well plate for 20 minutes. After incubation, cells were washed twice by centrifugation at 400 x g for 5 minutes using wash buffer comprising 2 mM EDTA and 1.5% FBS in 1× PBS to remove unbound antibodies. After washing, cells were analyzed using cytosoft software (Guava EasyCyte, Millipore, Billerica, MA). The percentage of live CD4^+^ and CD8^+^ T-cells was analyzed after gating on live cells based on forward-scatter and side-scatter.

### Nitrite production from adherent splenic and cecal tonsil mononuclear cells of chickens

On d 18 and d 28 post-hatches, single cell suspensions of cecal tonsils and spleen mononuclear cells were obtained as described earlier [16]. Nitrite production from adherent cells was carried out according to a previous method [17] with some modifications. Briefly, mononuclear cells were seeded at 1 × 10^6^ splenocytes/well in 96-well plates incubated for 24 hours at 37 °C in the presence of 5 % CO_2_, to allow them attach. Unattached cells were removed with media and replaced with RPMI freshly prepared as stated above. Adherent cells were stimulated with approximately 0 μg (200 μl RPMI-1640 culture media only) or 25 μg, 50 μg or 100 μg of ECP or 0.1 μg of *Salmonella Enteritidis* lipopolysaccharide (LPS) in 200 μl of RPMI-1640 culture media per well and incubated for 5 d at 37 °C in the presence of 5 % CO_2_. Samples were centrifuged at 630 X g for 5 min at 10°C and 100 μL of the supernatants were removed. The nitrite content of the supernatant was determined using a sulfanilamide/N-(1-naphthyl) ethylenediamine dihydrochloride solution (Ricca Chemical Co., Arlington, TX) following the manufacturer’s instructions. Nitrite concentrations were determined from a standard curve with sodium nitrite standards.

### Anti-ECP- and anti-flagellar-specific IgG and IgA antibodies in serum and bile of chickens

Serum from the and bile were collected from one bird per cage on d 18, and 2 birds per cage on d 21 and d 28 post-hatch. The amounts of anti-ECP- and anti-flagellar-specific IgG and IgA antibodies in serum and bile were determined by ELISA as previously described [11] with modifications. Native ECP was coated at 10 μg/ml (IgA) or 20μg/ml (IgG) on ELISA plates (Nunc Maxisorp™, ThermoFisher Scientific, Waltham, MA). Bile was diluted to 1:200 and serum was diluted to 1:20 in PBS containing 2.5 %, non-fat dry milk and 0.1% Tween 20 (VWR, Radnor, PA). Horseradish peroxidase (HRP) conjugated polyclonal goat anti-chicken IgG (Bethyl, Montgomery, TX) at 1:20,000 dilution or HRP-conjugated polyclonal goat anti-chicken IgA (SouthernBiotech, Birmingham, AL) at 1:10,000 was used as a secondary antibody. Optical density was measured as absorbance at 450nm using a spectrophotometer (Biochek, Scarborough, ME).

### Effect of serum antibodies from chickens on ECP neutralization

Liver male hepatoma cells (LMH) were seeded in 96-well plates at 4 × 10^4^ cells per well for 30 minutes in 100μl of Dulbecco’s modified eagle medium (DMEM). The DMEM was supplemented with 10% Fetal bovine serum and 1% Penicillin and Streptomycin. Antigen-antibody reaction was prepared in duplicates by incubating 15μl of *C. perfringens* supernatant containing 825μg protein with 15μl serum from each bird (n=6) for 60 minutes. The antigen-antibody mixture was added to the growing LMH culture. Cells were incubated in 5 % CO_2_ at 37°C for 8 hours. LMH cytotoxicity was measured according to the protocol outlined by OPS diagnostics (OPS diagnostics Lebanon, NJ) to measure the release of lactate dehydrogenase (LDH) from dying cells. Briefly, 50μl of cell-free supernatant of LMH culture was incubated with 50μl of 50mM Lithium Lactate and 50μl of a mixture composed of 100μl of 9mg/ml Phenazine methosulfate (PMS), 100μl of 33mg/ml iodonitrotetrazolium 2-(4-iodophenyl)-3-(4-nitrophenyl)-5-phenyl-2H-tetrazolium) (INT) and 2.3ml of 3.7mg/ml Nicotinamide adenine dinucleotide (NAD) solution. Color development was allowed for at least 5 minutes. Absorbance was measured at 490nm using a spectrophotometer (Biochek, Scarborough, ME).

### Effect of vaccination and challenge on the cecal tonsil mRNA levels of tight junction proteins

Total RNA from jejunum and ileum tonsils were collected from 2 birds per cage on d 21 and d 28 post-hatch. The mRNA was reverse transcribed into cDNA using methods described earlier [18]. The mRNA was then analyzed for the tight junction proteins claudin-2 and zonula occuloden-1 by real-time PCR using SyBr green, after normalizing for β-actin mRNA. Primer sequences and annealing temperatures are described in Table 1. The fold change with respect to the reference gene was calculated using the method 2^(Ct Sample−Housekeeping)^/2^(Ct Reference−Housekeeping)^ with Ct being the threshold cycle [19]. The Ct value was determined by CFX Maestro software when the fluorescence rises exponentially 2-fold above background (Biorad, Hercules, CA.). The reference group was the negative control birds.

**Table 1.**
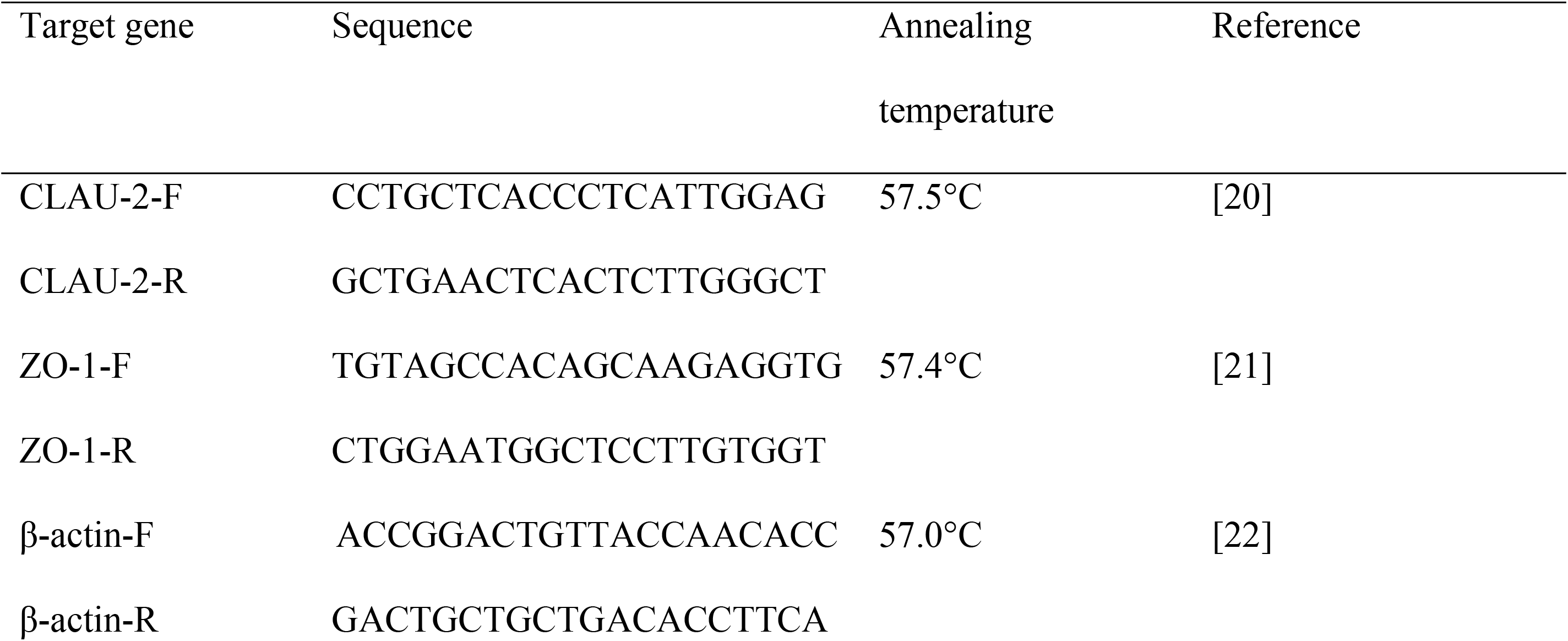
Primers and PCR Conditions for RT-qPCR.

### Statistical Analysis

All statistical analyses were carried out with statistical software (SAS, v. 9.4, SAS Institute Inc., Cary, NC, USA). Analysis of body weight gain (WG), feed conversion ratio (FCR), feed intake (FI), toxin neutralization, nitrite production, antibody and T lymphocyte responses was carried out using a one-way analysis of variance (ANOVA) and pre-planned orthogonal contrasts to compare negative control vs vaccinated-challenged, and vaccinated-challenged vs negative control. Analysis of mortality and gut lesions was carried using Kruskal-Wallis chi-square test. Significance was determined at P < 0.05 and/or at P < 0.01. P values less than 0.1 were determined as approaching significance.

## Results

### Effect of vaccination and challenge on mortality and jejunum lesions of broiler birds

Birds in the vaccinated-challenged group had comparable mortality to that in the negative control group, from d 14 to d 21 (one-week post challenge) and d 14 to d 28 post-hatch (two weeks post challenge), but had lower mortality than that in the positive control group which approached significance (P = 0.07, Table 1).

Birds in the vaccinated-challenged group had higher lesion scores (Table 2) than that in the negative control group but had numerically lower lesion scores than that in the positive control group which approached significance (P = 0.05, Table 2) on d 21 post-hatch.

### Effect of vaccination and challenge on feed intake, average body weight gain and feed conversion ratio of broiler birds

All three treatment groups had comparable FI, WG and FCR from d 0 to d 14 post-hatch (Table 3). However, the vaccinated-challenged group had numerically lower feed intake than that in the negative control group which approached significance (P = 0.08, Table 3), but had comparable feed intake to the positive control group from d 14 to d 21 post-hatch (Table 3).

The vaccinated-challenged group also had comparable FI to the negative control group (Table 3) but had statistically significantly higher FI than that in the positive control group from d 14 to d 28 post-hatch (p < 0.05, Table 3).

The vaccinated-challenged group had comparable FCR to the negative control group (Table 3) but had statistically significantly lower FCR than that in the positive control group from d 14 to d 21 post-hatch and from d 14 to d 28 post-hatch (p < 0.01, Table 3)

### Effect of vaccination and challenge on the proliferation of splenic PBMCs of broilers birds

Splenic mononuclear cells obtained from all 3 treatment groups on d 18 post-hatch and stimulated with 0 μg ECP, 50 μg ECP and 0.1 μg Con A had comparable proliferation (Fig 1A). Splenic mononuclear cells obtained from the vaccinated-challenged group on d 18 post-hatch and stimulated with 25 μg ECP had comparable proliferation to that in the negative control group (Fig 1A) but had lower proliferation than that in the positive control birds approaching significance (p = 0.09, Fig 1A). Splenic mononuclear cells obtained from the vaccinated-challenged group on d 18 post-hatch and stimulated with 100μg ECP had comparable proliferation to that in the negative control group (Fig 1A), but statistically significantly lower proliferation than that in the positive control group (p < 0.01, Fig 1A).

**Fig 1.**
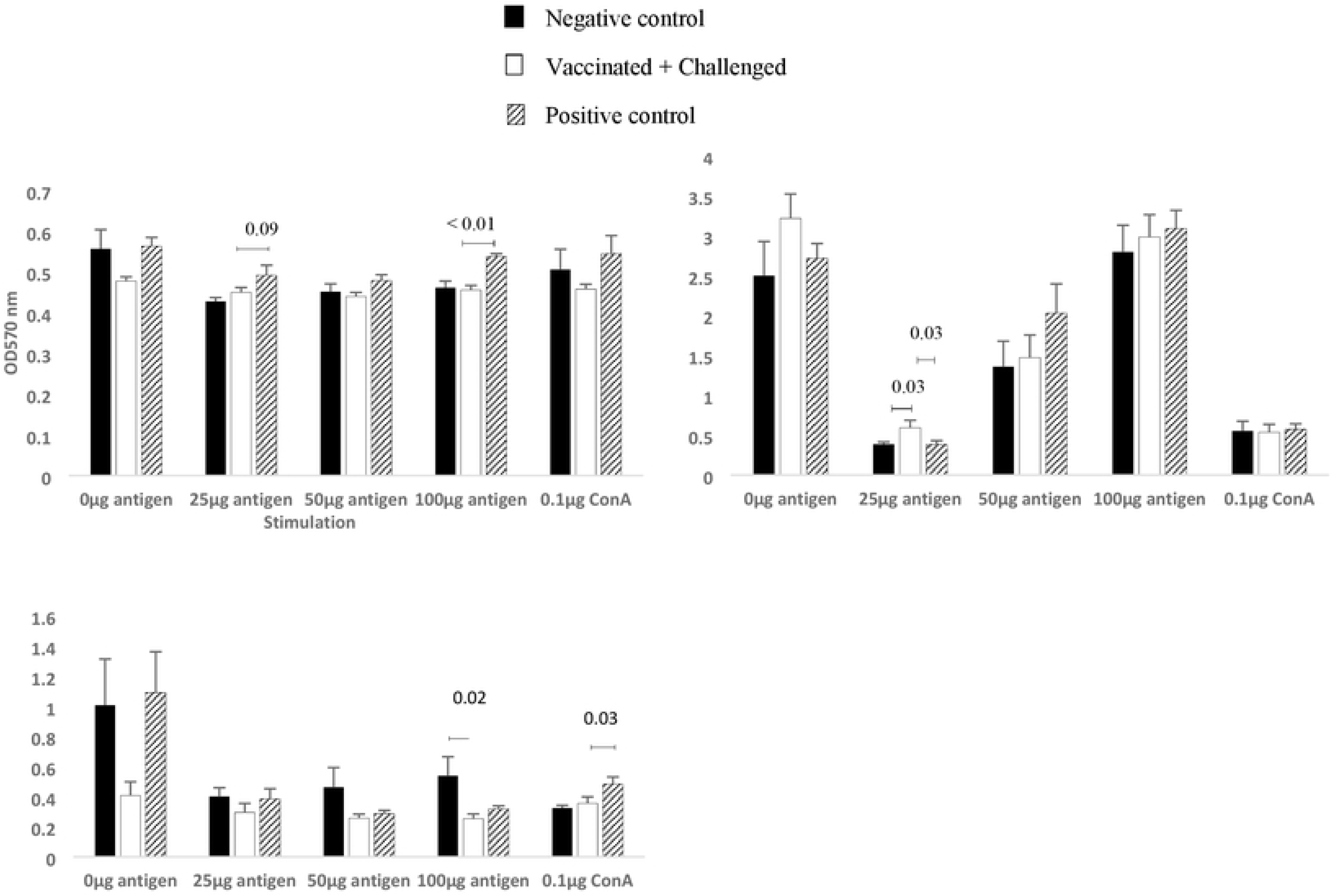
Effect of broiler vaccination and challenge on the proliferation of splenocyte mononuclear cells. Broiler chickens were orally vaccinated on day 0, day 4 and day 14 post-hatch with Chitosan-nanoparticles entrapped with *C. perfringens* ECP and *Salmonella* Enteritidis flagellar proteins, and challenged with *Eimeria* on day 14 and *C. perfringens* on day 19, 20, and 21 post-hatch. (A) On day 18, (B) day 21 and (C) day 28 post-hatch, splenic mononuclear cells were stimulated with 0.05, 0.1, 0.25 or 0.5 μg ECP for 5 days. Lymphocyte proliferation was measured using MTT assay and values reported as Optical Density (OD) values. Mean ± SEM of 6 replicates (n = 6). P < 0.05.

Splenic mononuclear cells obtained from all 3 treatment groups on d 21 post-hatch, and stimulated with 0μg ECP, 50μg ECP, 100μg ECP, and 0.1μg Con A had comparable proliferation (Fig 1B). Splenic mononuclear cells obtained from the vaccinated-challenged group on d 21 post-hatch and stimulated with 25μg ECP had statistically significantly higher proliferation than that in the negative control and positive control groups (p < 0.05, Fig 1B).

Splenic mononuclear cells obtained from all 3 treatment groups on d 28 post-hatch and stimulated with 0μg ECP, 25μg ECP, and 50μg ECP had comparable proliferation (Fig 1C). Splenic mononuclear cells obtained from chickens in the vaccinated-challenged group on d 28 post-hatch and stimulated with 100μg ECP had statistically significantly lower proliferation than that in the negative control group (p < 0.05, Fig 1C) but had comparable proliferation to the positive control group (Fig 1C). Splenic mononuclear cells obtained from chickens in the vaccinated-challenged group on d 28 post-hatch, and stimulated with 0.1μg ConA had comparable proliferation to that in the negative control group but had statistically significantly lower proliferation than that in the positive control group (p < 0.05, Fig 1C).

### Effect of broiler vaccination and challenge on the percentages of splenic and cecal tonsils CD4^+^, CD8^+^ cells, double positive CD4^+^ - CD8^+^ cells and CD4^+^:CD8^+^ratio

Splenic mononuclear cells obtained from all treatment groups on d 18, 21 and 28 post-hatch had comparable proportion of CD4^+^ cells (Fig 2A). Cecal tonsil mononuclear cells obtained from all treatment groups on d 18 and 28 post hatch also had comparable proportion of CD4^+^ cells (Fig 2A). Cecal tonsil mononuclear cells obtained from the vaccinated-challenge group on d 21 post-hatch had a comparable proportion of CD4^+^ cells to that in the negative control group (Fig 2A), but had a lower proportion of CD4^+^ cells than that in the positive control group, approaching statistical significance (p = 0.07, Fig 2A).

**Fig 2.**
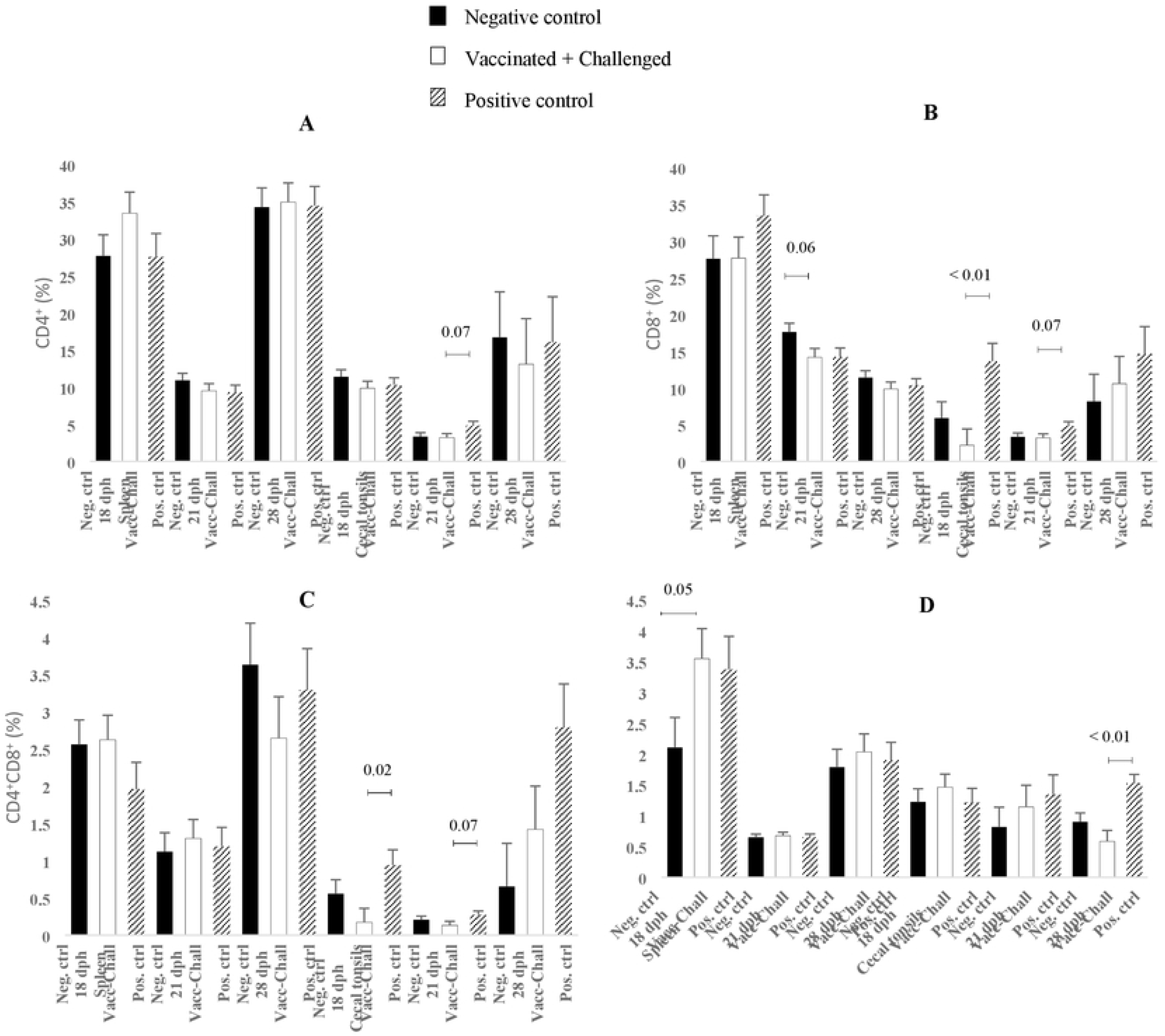
Effect of broiler vaccination and challenge on their CD4+ and CD8+ cell populations. Broiler chickens were orally vaccinated on d 0, 3 and 14 post-hatch with Chitosan-nanoparticles entrapped with *C. perfringens* ECP and *Salmonella* Enteritidis flagellar proteins, and challenged with *Eimeria* on day 14 and *C. perfringens* on day 19, 20, and 21 post-hatch. (A) Percentages of live cells that are CD4^+^ (B) Percentages of live cells that are CD8^+^ (C) Percentage of live cells that double positive CD4^+^ and CD8^+^. (D) The ratio of CD4^+^:CD8^+^ cells. Mean ± SEM. n = 6 replicates. P < 0.05.

Splenic mononuclear cells obtained from all treatment groups on d 18 and 28 post-hatch had a comparable proportion of CD8^+^ cells (Fig 2B). Splenic mononuclear cells obtained from the vaccinated-challenged group on d 21 post-hatch had a numerically lower proportion of CD8^+^ cells than that in the negative control group approaching statistical significance (p = 0.06, Fig 2B) but had a comparable proportion of CD8^+^ cells to that in the positive control group (Fig 2B). Cecal tonsil mononuclear cells obtained from the vaccinated-challenge group on d 18 post-hatch had a comparable proportion of CD8^+^ cells to that in the negative control group (Fig 2B), but had a statistically significantly lower proportion of CD8^+^ cells than that in the positive control group (p < 0.01, Fig 2B). Cecal tonsil mononuclear cells obtained from the vaccinated-challenge group on d 21 post-hatch had a comparable proportion of CD8^+^ cells to that in the negative control group but had a numerically lower proportion of CD8^+^ cells than that in the positive control group, approaching statistical significance (p = 0.07, Fig 2B). Cecal tonsil mononuclear cells obtained from all treatment groups on d 28 post-hatch had comparable percentages of CD8^+^ cells (Fig 2B).

Splenic mononuclear cells obtained from all treatment groups on d 18, 21 and 28 post-hatch had comparable percentages of double positive CD4^+^ CD8^+^ cells (Fig 2C). Cecal tonsil mononuclear cells obtained from the vaccinated-challenge group on d 18 post-hatch had a comparable proportion of double positive CD4^+^ CD8^+^ cells to that in the negative control group but had a statistically significantly lower proportion of double positive CD4^+^ CD8^+^ cells than that in the positive control group (p < 0.05, Fig 2C). Cecal tonsil mononuclear cells obtained from the vaccinated-challenge group on d 21 post-hatch had a comparable proportion of double positive CD4^+^ CD8^+^ cells to that in the negative control group but had a numerically lower proportion of double positive CD4^+^ CD8^+^ cells than that in the positive control group, approaching statistical significance (p = 0.07, Fig 2C). Cecal tonsil mononuclear cells obtained from all treatment groups on d 28 post-hatch a had comparable proportion of double positive CD4^+^ CD8^+^ cells (Fig 2C).

Splenic mononuclear cells obtained from the vaccinated-challenged group on d 18 post-hatch had a numerically higher CD4^+^:CD8^+^ ratio than that in the negative control group approaching significance (p = 0.05, Fig 2D) but had a comparable CD4^+^:CD8^+^ ratio to that in positive control group (Fig 2D). Splenic mononuclear cells obtained from all treatment groups on d 21 and d 28 post-hatch had comparable CD4^+^:CD8^+^ ratios (Fig 2D). Cecal tonsil mononuclear cells obtained from all three treatment groups on d 18 and 21 post-hatch had comparable CD4^+^:CD8^+^ ratios (Fig 2D). Cecal tonsil mononuclear cells obtained from the vaccinated-challenge group on d 28 post-hatch had comparable CD4^+^:CD8^+^ ratio to that in the negative control group but had a statistically significantly lower CD4^+^:CD8^+^ ratio than that in the positive control group (p < 0.01, Fig 2D).

### Effect of vaccination and challenge on nitrite production from cecal tonsil mononuclear cells of broiler birds on day 18 and 28 post-hatch

Cecal tonsil mononuclear cells obtained from chickens in all 3 treatment groups on d 18 post-hatch and stimulated with 0μg ECP, 25μg ECP, 50μg ECP, and 100μg ECP had comparable nitrite production (Fig 3A). Cecal tonsil mononuclear cells obtained from the vaccinated-challenged group on d 18 post-hatch, and stimulated with 0.1μg LPS had comparable nitrite production to that in the negative control group (Fig 3A), but had statistically significantly lower nitrite production than that in the positive control group (p < 0.05, Fig 3A).

**Fig 3.**
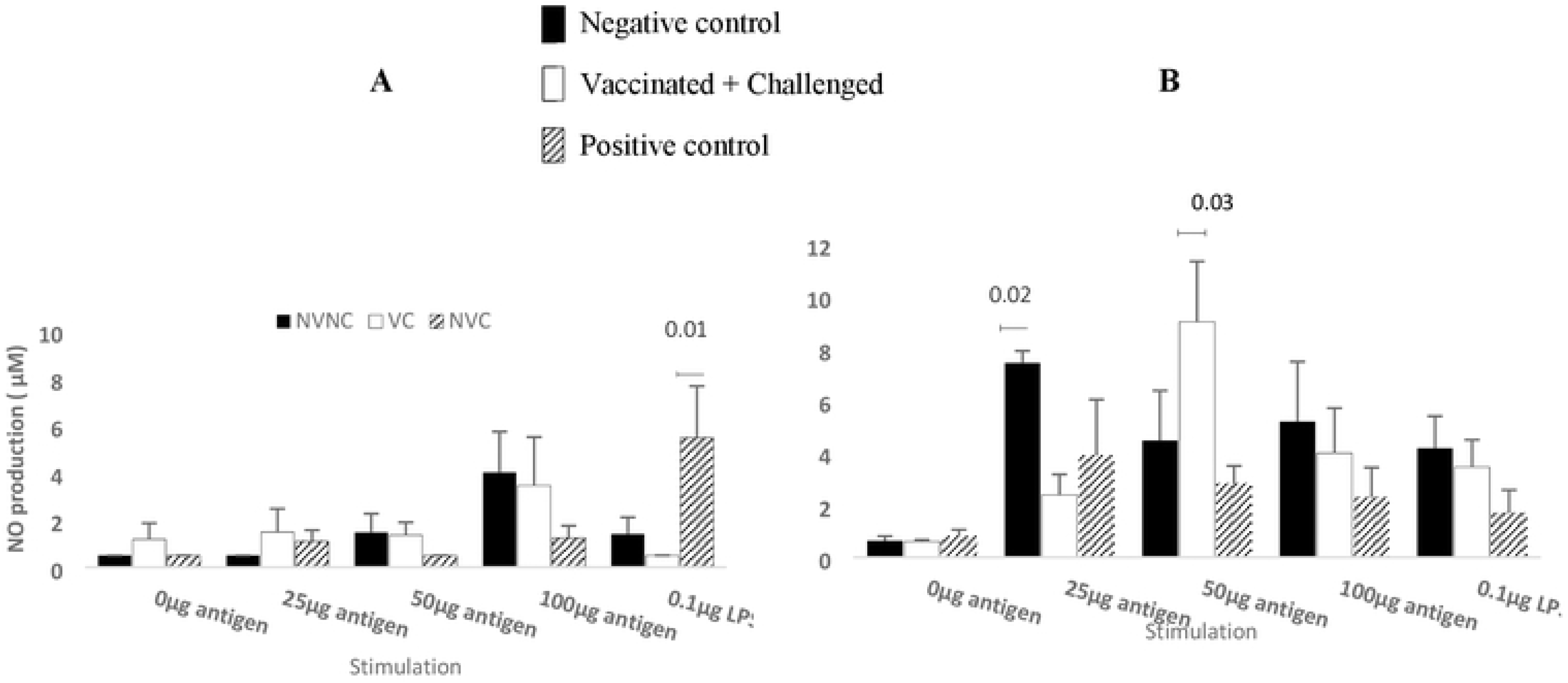
Effect of broiler vaccination and challenge on nitrite production from adherent cecal tonsil mononuclear cells. Broiler chickens were orally vaccinated on day 0, day 4 and day 14 post-hatch with Chitosan-nanoparticles entrapped with *C. perfringens* ECP and *Salmonella* Enteritidis flagellar proteins, and challenged with *Eimeria* on day 14 and *C. perfringens* on day 19, 20, and 21 post-hatches. On d 18 (Panel A), and d 28 (Panel B) nitrite production was measured using sulfanilamide/N-(1-naphthyl) ethylenediamine dihydrochloride solution. Nitrite concentrations were determined from a standard curve with sodium nitrite standards. Mean ± SEM of 6 replicates (n = 6). P < 0.05.

Cecal tonsil mononuclear cells obtained from all 3 treatment groups on d 28 post-hatch and stimulated with 0μg ECP, 100μg ECP and 0.1μg LPS had comparable proliferation (Fig 3B). Cecal tonsil mononuclear cells obtained from chickens in the vaccinated-challenged group on d 28 post-hatch and stimulated with 25μg ECP had statistically significantly lower nitrite production than that in the negative control group (p < 0.05, Fig 3B), but had comparable nitrite production to that in the positive control group (Fig 3B). Cecal tonsil mononuclear cells obtained from chickens in the vaccinated-challenged group on d 28 post-hatch, and stimulated with 50μg ECP had comparable nitrite production to that in the negative control group (Fig 3B) but had statistically significantly higher nitrite production than that in the positive control group (p < 0.05, Fig 3B).

### Effect of vaccination and challenge on nitrite production from splenic mononuclear cells of broiler birds on d 18 and d 28 post-hatch

Splenic mononuclear cells obtained from all 3 treatment groups on d 18 post-hatch and stimulated with 0μg ECP, 50μg ECP, 100μg ECP, and 0.1μg LPS had comparable nitrite production (Fig 4A).

**Fig 4.**
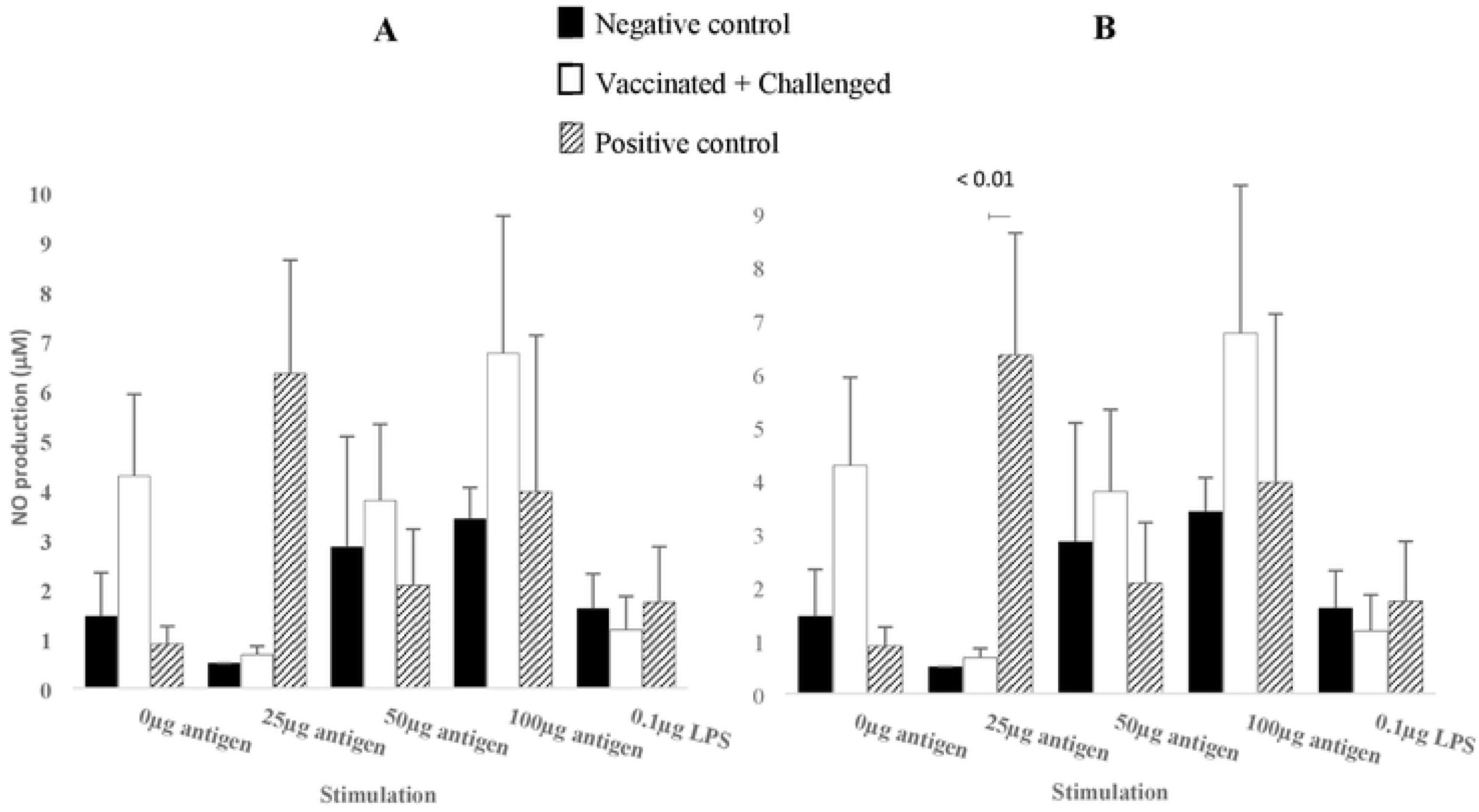
Effect of broiler vaccination and challenge on nitrite production from splenocyte mononuclear cells. Broiler chickens were orally vaccinated on day 0, day 4 and day 14 post-hatch with Chitosan-nanoparticles entrapped with *C. perfringens* ECP and *Salmonella* Enteritidis flagellar proteins, and challenged with *Eimeria* on day 14 and *C. perfringens* on day 19, 20, and 21 post-hatches. On d 18 and d 28 post-hatch, splenocyte mononuclear cells were stimulated with 0, 0.05, 0.1, 0.25 or 0.5 μg ECP for 48 hrs. (A) on day 18 and (B) on day 28 post-hatch, nitrite production was measured using sulfanilamide/N-(1-naphthyl) ethylenediamine dihydrochloride solution. Nitrite concentrations were determined from a standard curve with sodium nitrite standards. Mean ± SEM of 6 replicates. P < 0.05.

Splenic mononuclear cells obtained from all 3 treatment groups and stimulated with 0μg ECP, 50μg ECP, 100μg ECP, and 0.1μg LPS had comparable proliferation (Fig 4B). Splenic mononuclear cells obtained from chickens in the vaccinated-challenged group on d 28 post-hatch and stimulated with 25μg ECP had comparable nitrite production to that in the negative control group (Fig 4B), but had statistically significantly lower nitrite production than that in the positive control group (p < 0.05, Fig 4B).

### Effect of vaccination on anti-ECP-specific serum IgG and bile IgA

Serum and bile obtained on d 18, 21, and 28 post-hatches from all treatments had comparable anti-ECP antibodies in (Fig 5).

**Fig 5.**
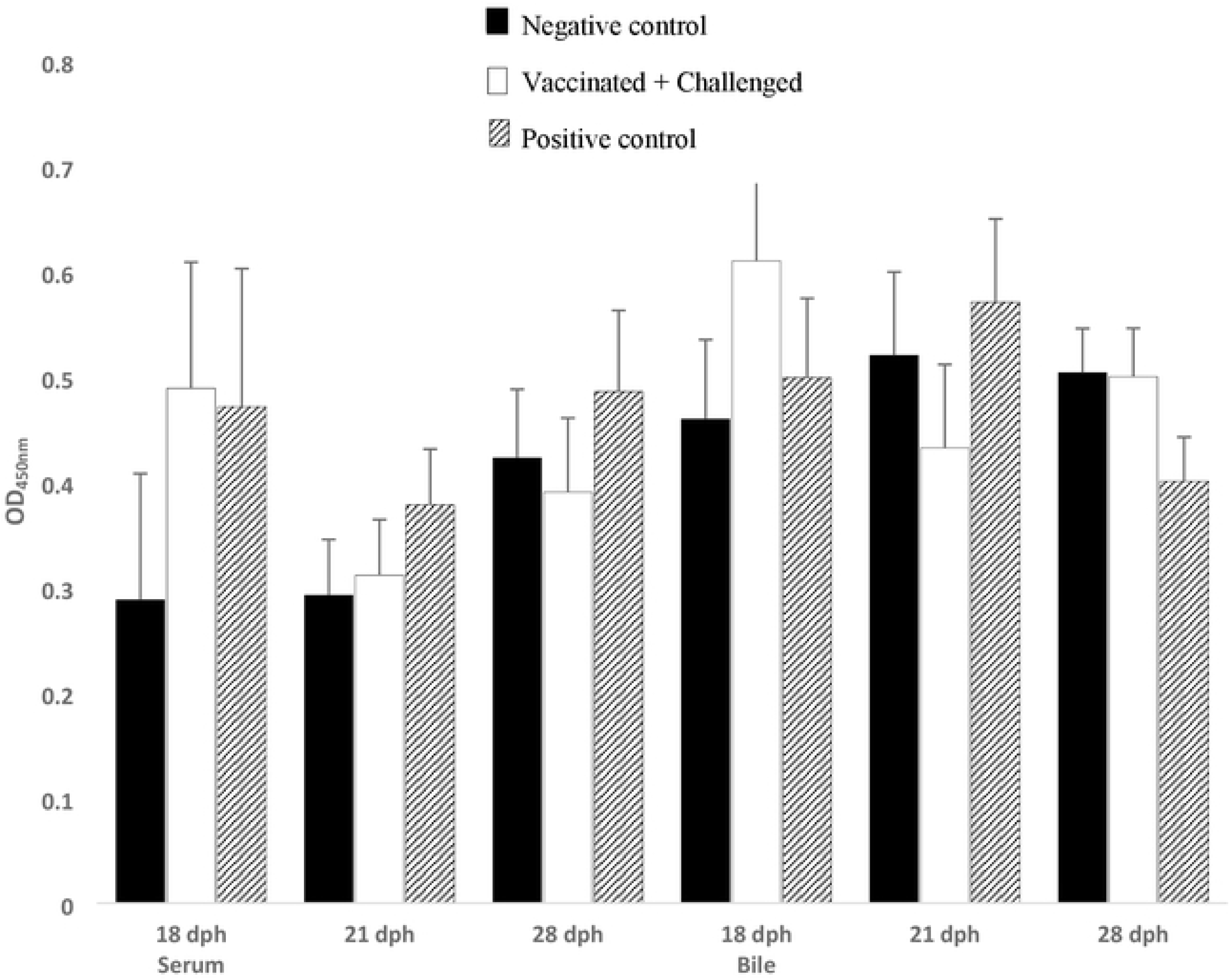
Anti-ECP-specific serum IgG and bile IgA. Broiler chickens were orally vaccinated on day 0, day 4 and day 14 post-hatch with Chitosan-nanoparticles entrapped with *C. perfringens* ECP and *Salmonella* Enteritidis flagellar proteins, and challenged with *Eimeria* on day 14 and *C. perfringens* on day 19, 20, and 21 post-hatch. Indirect ELISA was used to measure anti-ECP specific antibodies in serum and bile. Mean ± SEM of 6 replicates. P < 0.05.

### Effect of vaccination and challenge on ECP toxin neutralization by serum and bile antibodies

Serum obtained from all 3 treatments groups on d 18 and 28 post-hatch, and incubated with ECP and LMH cells had comparable LDH release (Fig 6A). Serum obtained from the vaccinated-challenged group on d 21 post-hatch and incubated with ECP and LMH cells had lower LDH release than that in the negative control group approaching statistical significance (p = 0.08, Fig 6A) but had statistically significantly higher LDH release than that in the positive control group (p < 0.05, Fig 6A).

**Fig 6.**
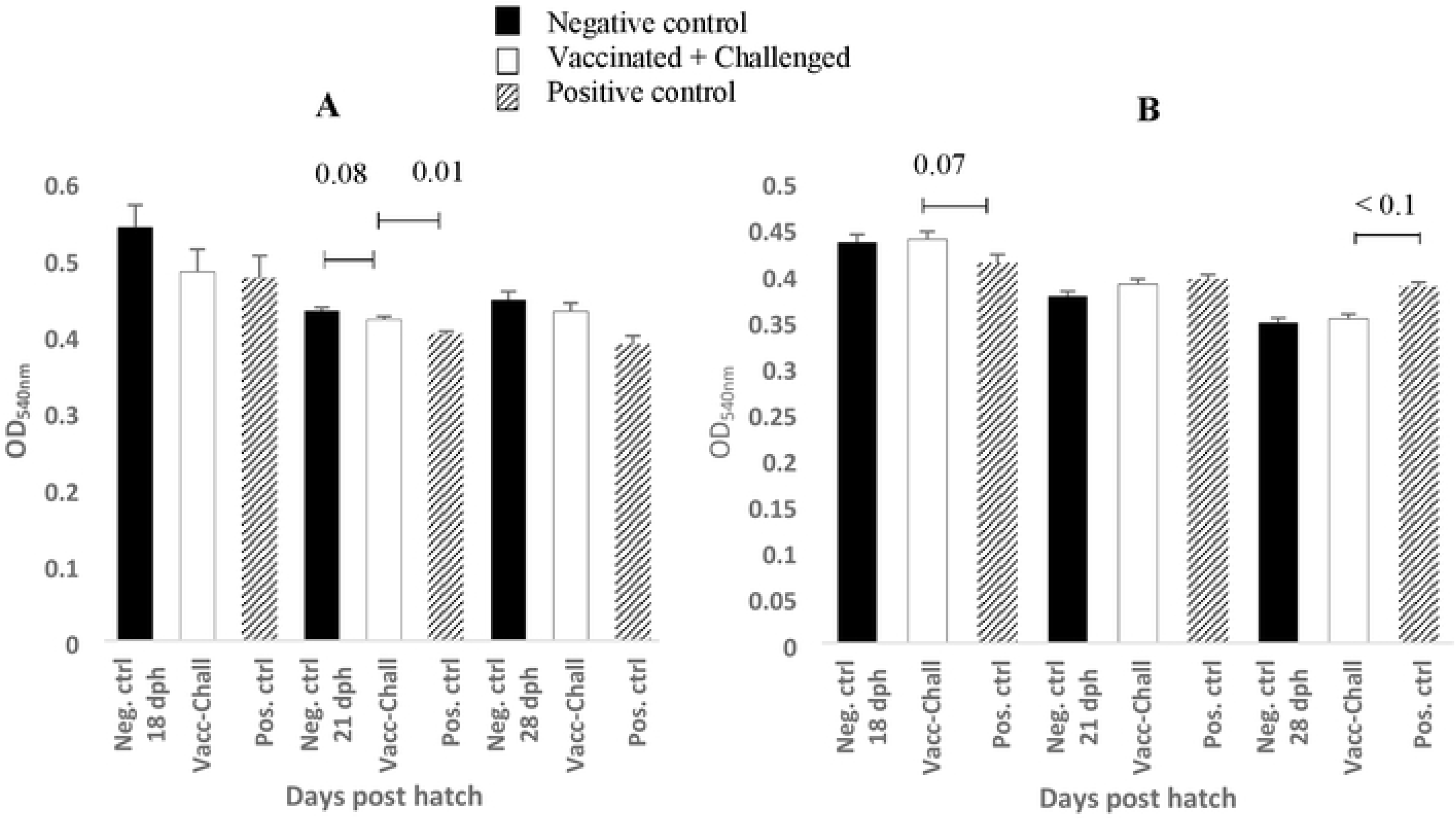
Effect of broiler vaccination and challenge on ECP toxin neutralization. Broiler chickens were orally vaccinated on day 0, day 4 and day 14 post-hatch with Chitosan-nanoparticles entrapped with *C. perfringens* ECP and *Salmonella* Enteritidis flagellar proteins, and challenged with *Eimeria* on day 14 and *C. perfringens* on day 19, 20, and 21 post-hatch. (A) serum and (B) bile were collected on day 18, 21, and d 28 post-hatch and incubated with 825 μg ECP to neutralize ECP. LMH cells were incubated with the above neutralizing solution for 8 hours. Cytotoxicity was measured by LDH release and values reported as Optical Density (OD) values. Mean ± SEM. n = 4 replicates. P < 0.05.

Bile obtained from the vaccinated-challenged group on d 18 post-hatch and incubated with ECP and LMH cells had comparable LDH release to that in the negative control group but had higher LDH release than that in the positive control group, approaching statistical significance (p = 0.07, Fig 6B). Bile obtained from all three treatment groups on d 21 post-hatch and incubated with ECP and LMH cells had comparable LDH release (Fig 6B). Bile obtained from the vaccinated-challenged group on d 28 post-hatch and incubated with ECP and LMH cells had comparable LDH release to that in the negative control group but had statistically significantly lower LDH release than that in the positive control group (p < 0.01, Fig 6B).

### Effect of vaccination and challenge on the jejunum and ileum mRNA levels of claudin-2, zonula occuloden-1, muc-2

Jejunum samples obtained from the vaccinated-challenged group on d 21 post-hatch had statistically significantly higher zonula occuloden-1 mRNA levels than that in the negative control and positive control group (P < 0.05, Fig 7). Ileum samples obtained from all treatment groups on d 21 post-hatch had comparable claudin-2, and zonula occuloden-1 mRNA levels (Fig 7).

**Fig 7.**
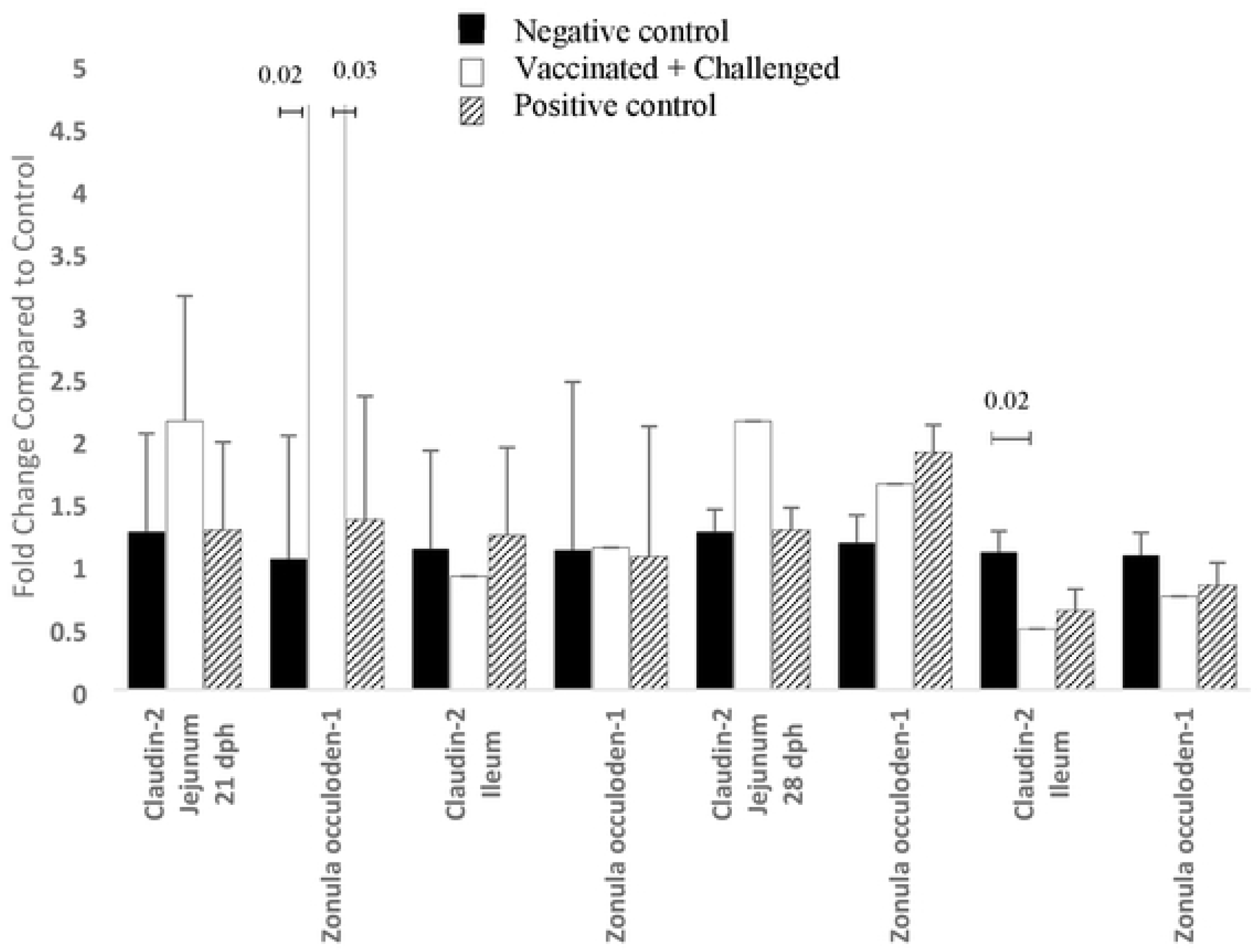
Effect of broiler vaccination and challenge on tight junction mRNA level. Broiler chickens were orally vaccinated on day 0, day 4 and day 14 post-hatch with Chitosan-nanoparticles entrapped with *C. perfringens* ECP and *Salmonella* Enteritidis flagellar proteins, and challenged with *Eimeria* on day 14 and *C. perfringens* on day 19, 20, and 21 post-hatch (dph). On day 21 and day 28 post-hatch, relative claudin-2 and zonula occuloden-1 mRNA content were analyzed after correcting for β-actin mRNA and normalizing to the mRNA content of the negative control group. Mean ± SEM. n = 6 replicates. P < 0.05.

Jejunum samples obtained from all treatment groups on d 28 post-hatch had comparable claudin-2 and zonula occuloden-1 (Fig 7). Ileum samples obtained from the vaccinated-challenged group on d 28 post-hatch had statistically significantly lower claudin-2 mRNA level than that in the negative control group (P < 0.05, Fig 7) but had comparable claudin-2 to that in the positive control group (Fig 7).

## Discussion

To the best of our knowledge, this study is the first to identify the immunogenicity and protective efficacy of nanoparticles loaded with *C. perfringens* ECP and *Salmonella* flagella proteins. The study was designed on the back of promising results from a previous study carried out by our lab that demonstrated the safety and immunogenicity of the same chitosan nanoparticle vaccine when administered by oral gavage to broilers, in the absence of an experimental challenge [15]. The experimental NE challenge model employed was successful in reproducing clinical NE as shown by 33% mortality and 0.72 average NE gut lesion in the unvaccinated challenged group. Over the 2-week period post challenge, the vaccinated-challenged group had numerically higher mortality and intestinal lesions than the negative control group but had numerically lower mortality and intestinal lesions than positive control birds. Vaccination also improved the feed conversion ratio and feed intake of challenged birds. Increased mortality, feed intake, lower feed conversion, body weight gain and the presence of intestinal lesions have been associated with clinical NE in broilers [23]. Vaccination did not significantly improve body weight gain of challenged birds at the intervals examined. However, the improvement in the other performance parameters suggests that the vaccine is partially protective against NE in broilers. In support of this, Hu et al. in 2013 [24] demonstrated that nanoparticle detained toxins are immunogenic and protective in mice. In addition, Zhao et al. in 2012 [13] demonstrated that chitosan nanoparticles vaccines loaded with Newcastle disease virus antigens offered protection against an oral challenge of Newcastle disease.

It has been shown that a cell-mediated immune response is more important for immunity to *Eimeria* infections [25] and chitosan has shown potential as an adjuvant to drive cell-mediated immunity [26]. It is not clear why splenic mononuclear cells from positive control birds had higher recall response than splenocytes from vaccinated-challenged and negative control birds on day 18 post-hatch, although higher proliferation was found in vaccinated-challenged birds compared to controls at the peak of infection, on day 21 post-hatch. A study by Zhao et al. in 2012 [13] also found increased lymphoproliferation of splenocytes of chickens vaccinated with chitosan nanoparticles encapsulating Newcastle disease virus compared to non-vaccinated controls. ConA is a non-specific T cell mitogen from plants that can be used to measure T cell proliferation in response to vaccination [27]. However, there was no increase in T cell proliferation in vaccinated-challenged birds suggesting either a different dose of ConA may be appropriate for ex vivo stimulation or vaccination was inducing a diminished T cell response at the assay time points. Furthermore, the phenotype of proliferating T cells can variably be dominated by effector or regulatory cells [28]. *Eimeria* has previously been demonstrated to induce Il-10 which can increase regulatory T cells [29]. Overall, there appears to be an antigen concentration-dependent effect on splenocyte proliferation. The decrease in proliferation of splenic mononuclear cells at 100μg ECP concentration on d 28 post-hatch may be due to the cytotoxicity of the ECP to a population of cells present at this time point [15].

The overall higher proportion of CD4^+^, CD8^+^, and double positive CD4^+^ CD8^+^ cells in the positive control birds compared to vaccinated challenge and negative control birds may explain the increased proliferation found in positive control birds discussed previously. According to Luhtala et al. in 1997 [30], unlike thymic CD4^+^ CD8^+^ double positive T-cells which are immature, peripheral CD4^+^ CD8^+^ double positive T-cells can respond to in vitro stimulation with antigens. Necrotic enteritis was also found to increase CD4^+^ and CD8^+^ and double positive CD4^+^ CD8^+^ cell population in chickens by Runhke et al. in 2017 [31]. A study by Hong et al. in 2006 [32] also demonstrated the persistence of relatively higher numbers of CD4^+^ T-cells during a secondary or challenge infection with *Eimeria*. The enhanced cell-mediated immune response associated with the positive control group may also be associated with increased disease pathology associated with susceptible chickens [9]. The numerical increase in the CD4:CD8 ratio on day 18 may be indicative of a Th2 response that generally results in antibody production [33] although there were no differences between treatments in the specific antibodies against ECP. There was an increase in the CD4:CD8 T cell ratio on day 28 post-hatch, in the cecal tonsils in positive control birds compared to vaccinated-challenged birds. The implication for this is not clear as the mechanisms underlying the temporal and spatial CD4 and CD8 response during NE is not yet fully understood.

Ex vivo stimulation of cecal tonsil mononuclear cells for nitrite production was carried out to assess innate priming. Vaccination resulted in an antigen-dependent modulation of nitrite production from ex-vivo stimulated cecal tonsil cells. LPS stimulated increased NO production in the positive control birds on day 18 post-hatch in the cecal tonsils, but down regulated NO production in the cecal tonsils by day 28 post-hatch. Nitric oxide is usually induced early during an infection and then rapidly cleared; else it can have a negative impact on bird performance [34]. iNOS has been demonstrated to be upregulated in chicken intestinal cells in response to *Eimeria* infection [35] but unchanged or down-regulated in chickens when *Eimeria* was co-infected with *C. perfringens*. The positive control birds compared to vaccinated and control birds, had increased NO production from ex vivo stimulated spleen mononuclear cells on d 28 post-hatch.

Indirect ELISA carried out to detect specific antibodies against ECP in serum or bile did not detect any differences between treatments. This may be because a crude antigen was used. Metabolic enzymes such as fructose 1,6 biphosphate aldolase and glyceraldehyde-3-phosphate dehydrogenase [36, 37] which are found in commensal organisms can generate antibodies against Clostridial infections which can interfere with ELISA results. Antigen neutralization assay was therefore carried out to determine whether undetected antibodies in serum or bile can neutralize toxins in ECP, thereby protecting LMH cells from lactose dehydrogenase release. LMH cells have been demonstrated to be vulnerable to LDH release by toxins such as NetB in the supernatant of virulent *C. perfringens* [38] and the strain of *C. perfringens* used for this study is *netB* positive (data not shown). Contrary to expectations however, increased cytotoxicity or decreased neutralization was observed in the serum and bile of vaccinated-challenged birds, compared to positive control birds on d 18 and d 21 post-hatch. The reason for this trend is not immediately clear, although for this study, endogenous complement was not inactivated and exogenous complemented was not added to serum or bile. Antibody-dependent and independent complement system has been characterized in poultry such as duck [39] and demonstrated to play a role *in vitro* in pathogen neutralization [40]. Also, non-specific complement cytotoxicity to certain cell lines is possible [41].

Quantitative PCR was carried out to determine the effect of vaccination on the mRNA levels of tight junction proteins such as claudin-2 and zonula occulodens-1. Claudin-2 is normally a pore forming protein associated with paracellular water and ion channels [42] while ZO-1 is positively associated with the formation of epithelial tight junctions by acting as tether for claudins [43]. A study by Bortoluzzi et al. in 2019 [44] showed that necrotic enteritis upregulates claudin-2 and downregulates ZO-1 at 21 days post-hatch. The decreased expression of claudin-2 in vaccinated-challenged and positive control birds compared to negative control birds on d 28 post hatch may therefore be associated with recovery. However, there was no difference between vaccinated-challenged and positive control birds claudin-2 levels.

Vaccination improved the expression of ZO-1 compared to negative control and positive control birds and this may have contributed to the improved performance of vaccinated-challenged birds compared to positive control birds.

## Conclusion

Chitosan nanoparticles loaded with ECP of *C. perfringens* and *Salmonella* flagellar proteins were immungenic and partially protective agianst experimentally induce NE. The vaccine produced cell mediated and humoral immune response in the birds. In order to understand the mechanisms underlying protection, further studies are required to identify the different populations of proliferating cells in vaccinted challenged, and non vaccinated challenged birds. Further studies are also needed to understand at the proteomic level, the expression of Th1 and Th2 cytokines. The vaccine was designed with only *C. perfringens* proteins with a homologous challenge. Further studies may be needed to determine heterologous protection, and if protection can be improved by adding *Eimeria* proteins, varying the route of adminstration, and refining the antigen selection.

## Acknowledgements

We acknowledge the efforts of the staff of Southern Poultry Research farms at which the field study was conducted. We also appreciate the support of members of the Selvaraj lab: Dr Theros Ng, Dr Jarred Oxford, Keila Acevedo, and Ragini Reddyvary for their contributing efforts during sampling collections and laboratory analysis.

